# Petal to the metal: The slow road to automating large-scale phenology labeling for herbarium specimens

**DOI:** 10.64898/2025.12.19.695552

**Authors:** Erin L. Grady, Raphael LaFrance, Daijiang Li, Russell Dinnage, Ellen G. Denny, John Deck, Robert P. Guralnick

## Abstract

Herbarium specimens represent critical historical records of plant phenology, yet automating annotation of reproductive structures remains challenging given the diversity of floral morphologies, specimen age and quality, and image quality. Here, we present a machine learning pipeline that uses an ensemble modeling approach to detect flowers on herbarium specimens and deliver these data to the phenology research community. After testing multiple strategies for generating training data, we found in-house expert-curated annotations were essential for producing reliable results. Expert validation found relatively strong accuracy for detecting present floral structures, but still had moderately high false negative rates. Applying the ensemble to our filtered final image dataset of 22 million records resulted in 11.1 million records labeled with flowers present. However, only 2.9 million of these contained complete metadata necessary for downstream phenology research, highlighting the need for full label digitization efforts. Still, this dataset represents a large compilation of historical herbarium-derived phenology records available as a resource for the phenology community. We end by demonstrating how integrating these machine-labeled records into Phenobase, a publicly-available phenology database, expands taxonomic and temporal coverage for large-scale phenological analyses, and discuss remaining challenges and next steps.

## INTRODUCTION

Natural life cycle events, such as plant flowering, fruiting, and leaf-out, occur annually on seasonally-defined schedules shaped by environmental cues. The timing and sequence of these events, which vary depending on species and environment, underpin many biological and human processes, including species interactions (Kudo et al., 2004), carbon cycling (Richardson et al., 2013), resource acquisition (Deacy et al., 2017), agriculture (Yadav et al., 2023), tourism (Nagai et al., 2019), and allergy management (Manangan et al., 2023). Because these seasonal changes are critical and fundamentally linked with many socio-ecological processes, a whole branch of inquiry, phenology, is devoted to understanding their drivers and consequences.

A fundamental question in phenology is how different species and ecosystems have responded and will continue to respond to accelerating human-driven environmental change. The rate and direction of these phenological shifts are of particular interest as they are often considered one of the first and best indicators of broader ecological disruption (Menzel et al., 2006). For example, such shifts can interfere with key interactions that require phenological synchrony, such as pollination by interacting insects, and can have cascading consequences for species’ population health, ecosystems, and people (Visser et al., 2019). Thus, access to data capturing the timing of key life stages across time, space, and taxa is increasingly of interest and importance across many scientific and applied disciplines.

Concerted phenology monitoring has been a key means to generate data needed to document phenological change. However, many regions across the globe don’t have phenology monitoring programs, and in areas that do, the temporal extent is often limited. For example, the USA National Phenology Network began coalescing observational monitoring in 2007 (Crimmins et al., 2022), but national-scale phenology data before this date are limited and often of short duration (with some exceptions, see for example, Hough’s efforts from 1851-1859; Guralnick et al., 2025). Beyond direct phenology monitoring, there are new resources, such as iNaturalist, producing digital vouchers where phenological information can be secondarily derived (Dinnage et al., 2025). Such contemporary contributory science platforms, like iNaturalist, harness community members across the globe to generate large quantities of observations linked to images and now include means to annotate phenological states. While these resources are helping to further fill spatial and taxonomic data gaps and providing new opportunities for phenology research at scale, they are even more limited in temporal extent compared to monitoring data.

A critical component for defining and predicting phenological change is understanding phenology and its trends over decadal to century-level timescales. Herbaria, libraries that house collections of pressed plant vouchers from the field, contain specimens dating back sometimes hundreds of years and have the potential to offer valuable historical phenology data, complementing contemporary monitoring initiatives (Davis et al., 2015; Willis et al., 2017). Recent efforts to image and digitize herbarium specimens have allowed for broader use of these plant collections (Soltis 2017), with information on over 108 million angiosperm specimens (∼40 million with associated images) available on the Global Biodiversity Information Facility (GBIF) as of October 2025. Ideally, each of these herbarium specimens contains valuable information about the location, collection date, species identity, and occasionally the phenological phase of that plant individual. For the vast majority of plants lacking phenology annotations, it requires significant effort to classify which, if any, phenological phases are visibly present on specimens.

Automating phenology annotation for digitized herbarium specimens is a clear next step, one that would unlock vast amounts of historical data and enable phenological change research across diverse taxa and environments. Recent machine learning efforts to classify phenological phases in photos of live plants have been successful (Dinnage et al., 2025); however, herbarium specimens present a unique set of challenges. Dried and pressed plant specimens often lose color and structure over time, reproductive structures can be lost during processing, some taxa have structures that are undetectable from digitized images alone or without dissection, and specimens contain non-plant material such as labels and herbarium stamps that may interfere with machine detection (Pearson et al., 2020). Additionally, accuracy of machine phenology annotation is highly dependent on the annotator who creates training data used to calibrate models, and curating a reliable and taxonomically-diverse training dataset requires extensive effort and careful consideration.

Several works have applied machine learning to herbarium specimens, with greater success in species identification than phenological recognition tasks (Hussein et al., 2022). Phenological efforts have often focused on small, curated species and image sets (Goëau et al., 2020; Younis et al., 2020) and many have reported challenges annotating specific phenological structures such as flowers (Lorieul et al., 2019; Younis et al., 2020). More recent efforts have successfully addressed specific phenology research questions using models trained by compositing many datasets from multiple different sources (Williamson et al., 2025). Our aim here, however, was not to produce a standalone research product or answer a particular question, but to develop a trustable, usable, and integratable data resource for the broader research community. Doing so required careful curation of our own training data, which allowed us to leverage information gathered during training to reduce noise in the final data product, as we discuss belowt.

Here, we present an automated phenology annotation pipeline that accurately detects flowers present on digitized herbarium specimens. With the goal of maximizing geographic and taxonomic coverage across the angiosperm tree of life, this effort unlocks vast quantities of previously limited historical phenology information. We outline the challenges, successes, complexities and strategic decisions involved in developing the pipeline, emphasizing the need for high-quality training data generated by botanically-knowledgeable individuals and rigorous filtering on multiple dimensions both pre- and post-modeling. We discuss the trade off between strict data filtering for improved model accuracy and data retention to maximize the volume of downstream historical records annotated with phenology information. We highlight a surprising and disappointing discovery that the majority of specimens lack metadata sufficient for phenological annotation, which we discovered after all the rest of the effort. Finally, we demonstrate the vast scale of previously-unavailable historical phenology data that are now available to stakeholders, and discuss the integration of these data into Phenobase (https://phenobase.netlify.app/), an online database of millions of research-ready, community-sourced phenology data from field images and from structured in situ phenology monitoring programs.

## METHODS

### Gathering specimen image and metadata

We opted to gather data for this work from GBIF, which provides access to billions of species occurrence records and, in many cases, associated images. We searched GBIF in mid-2023 for all “Magnoliopsida” and “Liliopsida”, using filters to select the basis of record as “preserved specimen” and media type as “images”. This produced 33,691,939 occurrence records with 35,314,854 corresponding multimedia records. We did not filter out records missing coordinates or location information so that when metadata of a record is updated in the future, its associated phenological information will have already been extracted. We downloaded this dataset and associated images in October 2024 and, using a nearest pixel filter, programmatically resized each image to 1024 pixels along the shorter edge to reduce storage and satisfy GPU memory limitations. Because services are often down and media links can change, the number of images downloaded is a subset of the total number of records in the original GBIF dataset. In sum, after contending with link issues, removing small or low quality images, and images associated with taxa that we decided to filter out (described below), we collected 22,447,563 images for downstream labeling and model training.

### Labeling training data

A challenging aspect of developing a reliable automated pipeline for phenophase annotation on herbarium specimens across the angiosperm tree of life was generating enough high-quality training data. Our goal was to produce a trusted historical phenology data resource for broad use, and we therefore deliberately chose to manually generate our own training data rather than utilizing data from multiple external sources. As mentioned above, manually scoring pressed plant specimens for phenophases is time consuming and difficult, and a key challenge is balancing the need for botanical expertise with the capacity of expert annotators. Other initiatives have employed volunteers, undergraduate students, and paid non-experts with relative success (Brenskelle et al., 2020; Zhou et al., 2018), however these focused on a few key taxa with easily-distinguishable structures. Limiting annotations to a few taxa at the genus or family level allows for more in-depth, targeted training and familiarity with target taxa which likely lead to better results. We were unsure whether employing non-experts would be successful when scoring thousands of diverse taxa.

We first tested whether a paid undergraduate student could produce accurate phenophase annotations across all flowering plants. We employed a student to score a random selection of specimens for presence or absence of open flowers. If the student was uncertain or unable to make an informed decision about the digitized image, they had the option to mark “unknown”. We then also had a botanically-knowledgeable individual validate these annotations to assess their accuracy. As reported in Results, we found the undergraduate’s open flower annotations were not at a level usable for training a model. Based on these results and other work that suggests that one of the most important variables in annotation accuracy is simply the annotator themselves (Brenskelle et al., 2020), we made the conscious choice to employ one in-house botanically-knowledgeable individual to generate all training data. This decision greatly reduced our capacity to generate high quantities of training data, but ensured high-quality and consistency, which was our priority.

In addition to the first set of specimens annotated to validate the undergraduate annotations, we annotated a second additional set of specimens for open flowers to increase training data. While compiling the second set, we randomly selected specimens in families underrepresented in the first set in an effort to promote taxonomic evenness. Additional records were added to the training data iteratively through repeated bouts of image validation throughout model training.

### Filtering training data

To ensure accurate and reliable results, we opted to filter out taxa pre-model-training that were challenging to manually annotate. First, we removed genera that had more than 5 records in the training dataset and an “unknown” annotation rate greater than or equal to 25%. After removing these genera, we removed families using the same criteria. Many of these removed taxa were those with small reproductive structures, such as Chenopodiaceae, or those with flowers and fruits that are difficult to distinguish from each other in images alone, like Poaceae. These taxa were removed permanently from our pipeline, both from training and downstream data. We made this choice strategically to balance data quality and quantity. Throughout this process, we found it critical to acknowledge and respond to limitations, both in human expertise, machine capabilities, and overall capacity. Final lists of removed genera and families are available in Supporting Information.

### Model training and post-training thresholding

#### Model Ensembling Approach

One challenge with training models on all herbarium data is the vast and complex floral diversity across the whole of the group, challenging models to properly abstract features that can apply broadly and that are not easily confusable with other objects. We realized single models often performed modestly well, but that ensembled models might perform better than any single one given this variability. We tried several different neural networks, architectures, and model parameters and ultimately determined that an ensemble modeling approach, combining three models, was most effective based on downstream validation statistics. Note that all three models were fine tuned versions of models that were previously pretrained on ImageNet data (Deng et al., 2009).

Despite differences across the three models (Effnet, ViT_MSE_ and VIT_CE_, summarized in Table 1), all training followed the same general steps. First, expert-labeled records were split into 60% training, 20% testing, and 20% validation sets. After low-quality images were filtered out, the images were resized to match the size requirements of the specific model (Table 1). Images in the training dataset were augmented using random horizontal and vertical flips and “auto augment” (Cubuk et al., 2018), which uses multiple techniques including random rotation, color jitter, sharpness adjustment, and image shear. Next, all models were color adjusted to use the ImageNet color mean and standard deviation. During training, each model was saved as to five checkpoints The final best model was chosen as the checkpoint with the best F1 score on the holdout validation dataset. The F1 score is the harmonic mean of precision (how many positive model predictions are actually positive?) and recall (how many actual positives did the model detect?). We chose the F1 score because it is particularly useful for assessing predictive performance for the positive class (“flowers present”), which is the focus of this effort. This overall approach is standard for model selection and ensures comparable scoring across model approaches (e.g. EffNet CNN versus ViT) and that models are tuned, selected, and ensembled based on their best validation performance.

**Table 1.** Description of the three models included in the ensemble modeling approach. Threshold values are cut-offs applied to each model’s continuous output probabilities. The difference between model 2 and model 3 is the loss function used. Mean square error loss functions are more akin to confidence scores and are typically Gaussian, while cross-entropy is better suited for classification tasks and tends to skew continuous probabilities more strongly to 0 or 1.

| Model type | Image size requirements | Threshold low | Threshold high | Loss function |
| --- | --- | --- | --- | --- |
| EffNet CNN | 528 x 528 pixels | 0.05 | 0.95 | Mean squared error |
| Visual transformer network (ViT) | 384 x 384 | 0.5 | 0.7 | Mean squared error |
| Visual transformer network (ViT) | 384 x 384 | 0.5 | 0.7 | Cross entropy |

We ran models using a single A100 GPU on the University of Florida’s HiPerGator. Each model was trained for at least 200 epochs, and we saved five checkpoints during training. At the end of every epoch, we evaluated model performance on a held-out validation set that was never used for backpropagation, which we used to assure model generalization and select the best checkpoint. We used several metrics for evaluating the model, including F1 score, accuracy, validation set loss, precision, and recall. After training, each checkpoint from all three models was post-processed to determine the probability thresholds that maximized accuracy. We divided the range [0.0–1.0] into three regions defined by two thresholds. Images with a score above the high threshold were considered a positive indication of the trait, images with scores below the low threshold were considered a negative indication of the trait. Images with scores between the high and low thresholds were classified as “equivocal”, meaning the model could not decide if the trait was present or absent. We adjusted those two thresholds until the images in the test (holdout) dataset accuracy was at the maximum. We also guarded against equivocal records being more than 30% of the test dataset. We used a batch size of 32, the initial learning rate was 5e-5 with a weight decay parameter of 0.01, and an AdamW optimizer.

For each image, each model generated a final outcome of positive, negative, or equivocal. To ensemble our three models, we used a two-thirds “majority rule” approach, focusing in particular on predictions of “flowers present” (e.g., where at least two models of the three predicted present with high certainty). Since herbarium specimens often contain only a portion of a plant individual and absence on the whole plant cannot be inferred, we focused on the accuracy of “present” labels and only retained images machine-labeled as “present” in downstream data. All other cases were scored “flowers not present” or “equivocal”. Note that the ensemble itself can be undecided (equivocal), for instance with [positive, negative, equivocal] or two or more equivocal votes from the three model outputs.

#### Model validation

We validated ensemble results both using held-out test data and through additional expert validation. For the expert validation, we machine-labeled a random subset of images and validated 600 specimens machine-labeled as “present” for flowers and 300 specimens machine-labeled as “absent” for flowers. Consistent with the Plant Phenology Ontology (Stucky et al., 2018), “flowers” here refers to any floral structure, including both flower buds and open flowers.

Although the models were originally trained on “open flowers”, we opted to conduct validation at the broader “flower” level, as these tasks are difficult and we felt “flower” better reflected the features detected by the ensemble models. To further understand instances where the model consensus was incorrect, we anecdotally looked at false positives and false negatives to understand general patterns in the mistakes (see Results).

### Machine labeling and integration with other sources

After model validation, the remaining unlabeled downloaded images were passed through the ensemble model pipeline. We retained only images labeled as “present” by at least two out of three models and data fields were standardized for integration in Phenobase, which has a defined set of required and optional fields. As we discuss below, we dropped records that we labelled but were missing these key fields that are required for phenology research. Key fields here include: scientific name, event date, and decimal latitude and longitude. The remaining data were made publicly available through the Phenobase web portal (https://phenobase.netlify.app/).

### Evaluating specimens missing critical fields

A key challenge and major limitation during this process was that a large number of specimens did not have date or location information included in their metadata. As these two pieces of information are critical for meaningful phenology data, this resulted in the loss of substantial data. To further investigate this issue, we randomly selected 100 specimens labeled as “flowers present” that were missing both date and location information. We manually annotated these specimens for whether or not they included specific (day, month and year) date information and georeferencable locality information on the specimen label.

## RESULTS

### Labeling training data

We worked with an undergraduate student to annotate a total of 3,525 specimens for open flowers, providing them with resources for the task and asking them to use online resources when needed. When validating these annotations, the undergraduate and one of our team members (lead author EG), who served as a botanical expert, were in agreement 69% of the time. Given this result, we chose to not include the undergraduate annotations in our final training dataset. EG then generated a total of 9,004 annotations from individual specimens for open flowers. After removing 49 genera and an additional seven families (Appendix S1; see Supporting Information with this article), 1660 genera and 261 families remained. Our final training dataset contained 7,205 specimen records, 4319 annotated as open flowers present and 2886 annotated as open flowers absent. Once expert-labeled records were split, 4324 images were included in the training set, 1442 in the testing set, and 1439 in the validation set.

### Model fitting and validation

We utilized three models (Table 1) and ensembled their results to predict the presence of flowers. Our training model accuracy scores across all three models varied between 88.3% and 92.5%. In general, the vision transformers models (ViT) performed better (92.2% and 92.5% accuracy) than the EffNet model (88.3% success). Overall, from the full ensemble, validation on the held-out test data found 96.3% accuracy for detecting the presence of flowers and 86.8% accuracy for detecting the absence of flowers (Appendix S2). However, expert validation on records not in the initial training, testing, or validation processes found the ensemble models to be 90.5% accurate for detecting the presence of flowers and 52.3% accurate for the absence of flowers (Table 2). The low accuracy for detecting absences was expected and likely due both to expert-validated data being more diverse than training data and that we were less concerned with training for absences in our overall process. Further investigation of the false positives from the expert validation found that the majority of false positives were fruits mistaken for flowers, although some sheets had no reproductive structures present and one sheet had a fruit illustration (Figure 1). Investigation into false negatives from the expert validation found a variety of reasons for the mistakes, many explainable, including situations where flowers weren’t attached to the main plant, flowers that may have been confused as fruits, and flowers that were very challenging to see (Figure 2).

**Figure 1.**
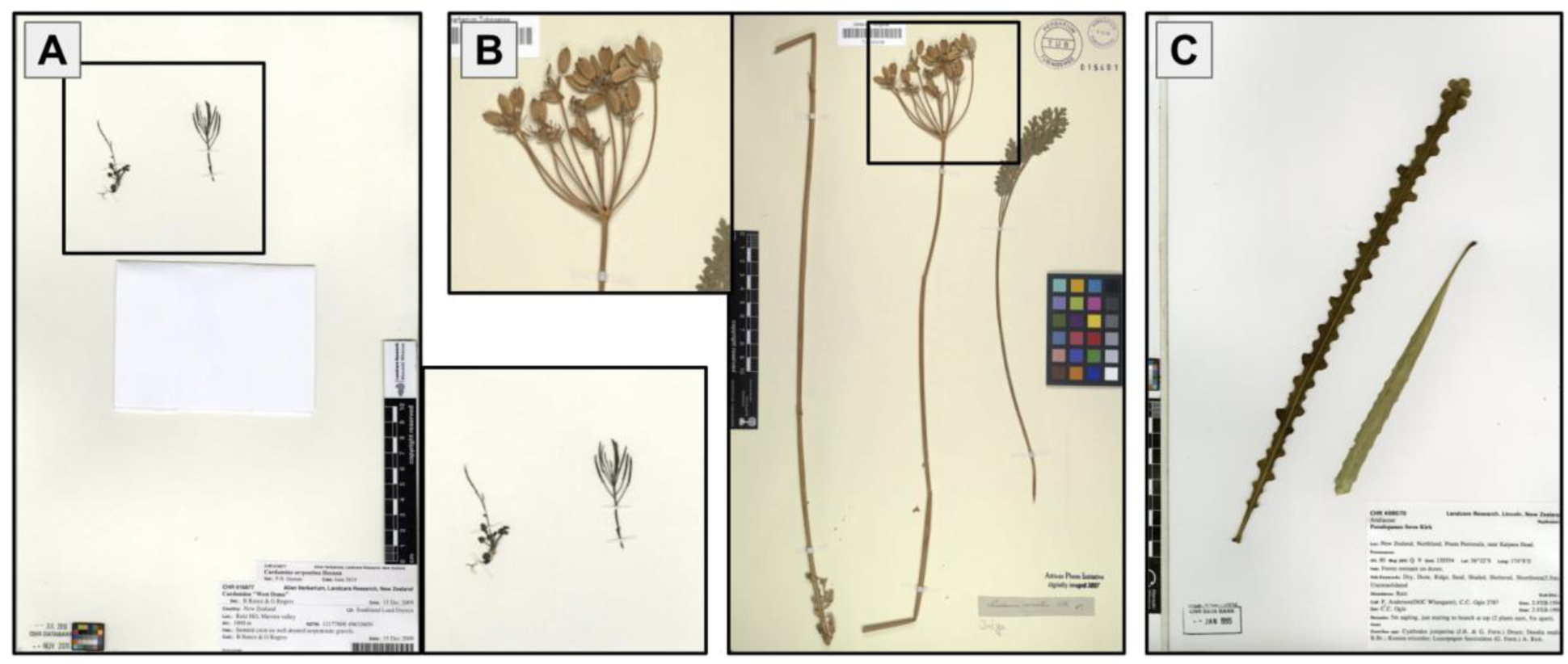
Examples of false positives, situations where the ensemble model consensus scored the image as “flowers present”, but there are no flowers present in the image. A. and B. are examples where fruits were confused for flowers. This was the most common cause of false positives, accounting for over half of cases. C. is an example where leaves were confused as flowers. Around 30% of false positives were situations where vegetative (non-reproductive) structures were confused as flowers.

**Figure 2.**
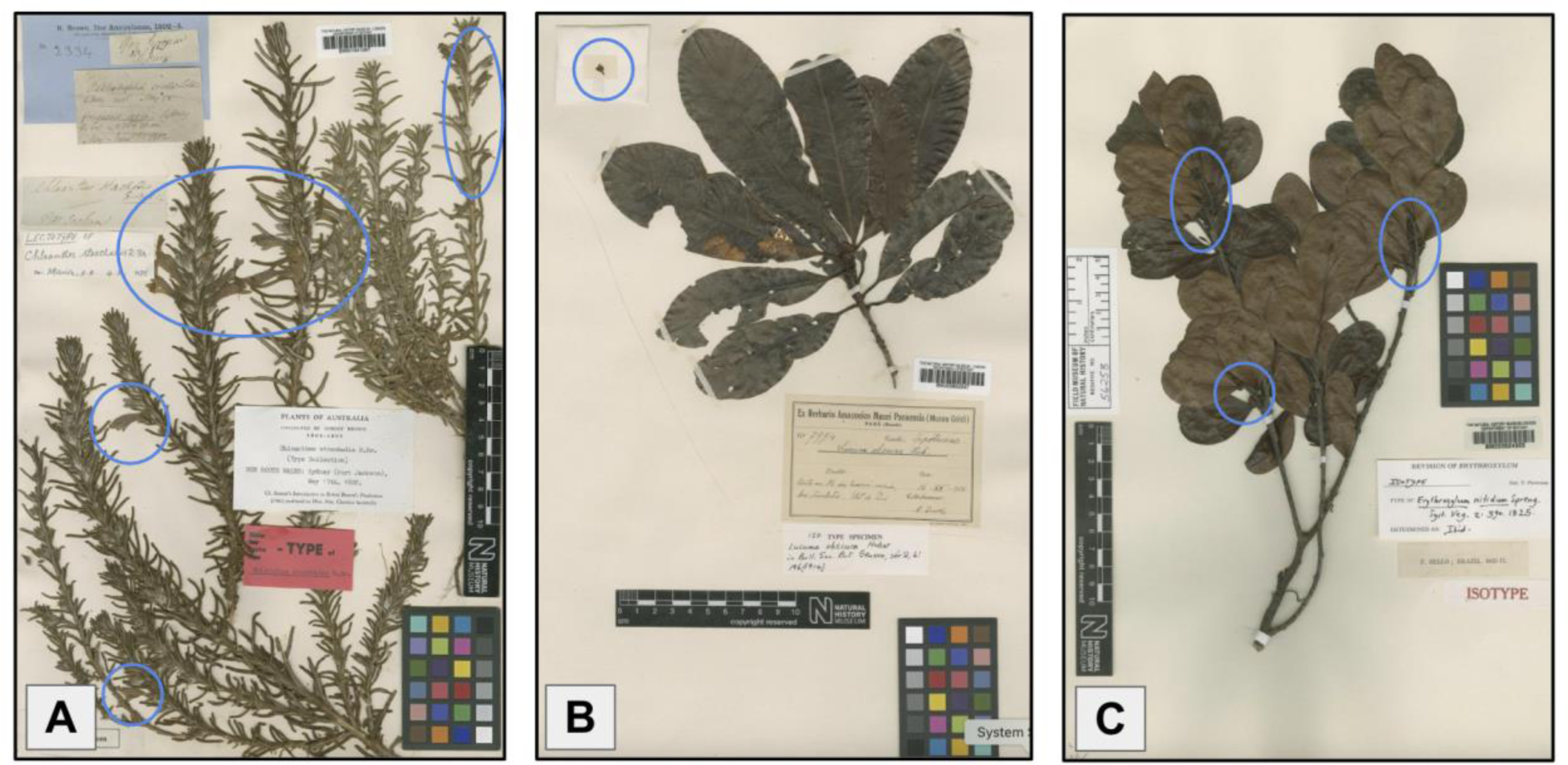
Examples of false negatives, situations where the ensemble model consensus scored the image as “flowers absent”, but there are flowers present in the image. A. is an example of a situation where there are obvious flowers on the sheet and the machine simply missed them. B. is an example of a situation where the floral structure is not connected to the plant. This often caused the machine to miss the flowers in the image. C. is an example where flowers are present, but just really challenging to see.

**Table 2.** Expert validation of machine-labels for flowers present or absent. Bolded values represent situations where the human expert and model consensus were in agreement. Grey cell represents the accuracy relevant to the final, downstream data.

| Expert Annotation | Model consensus: Present | Model consensus: Absent |
| --- | --- | --- |
| Present | **90.5% (543)** | 47.7% (143) |
| Absent | 9.9% (57) | **52.3% (157)** |

### Machine labeling and integration with other sources

After downloading and filtering out problematic images (images that could not be downloaded, images that cannot be read, or images that were either less than 10 KB or greater than 32 MB) and images of hard-to-annotate genera and families (Appendix S1), we were left with 22,447,563 images of digitized herbarium specimens. These images were fed through the ensemble modeling pipeline and annotated for presence or absence of flowers. Of these records, 11,131,820 were labeled as flowers present (meaning at least ⅔ models scored as present), but only 2,913,353 of these contained valid latitude, longitude, and date information (Figure 3). 5,748,788 images were scored as flowers absent (meaning at least ⅔ models scored as absent), and 5,566,955 were equivocal (meaning there was no consensus among the models). Only records scored as flowers present and with the needed metadata were retained and integrated into Phenobase (https://phenobase.netlify.app/).

**Figure 3.**
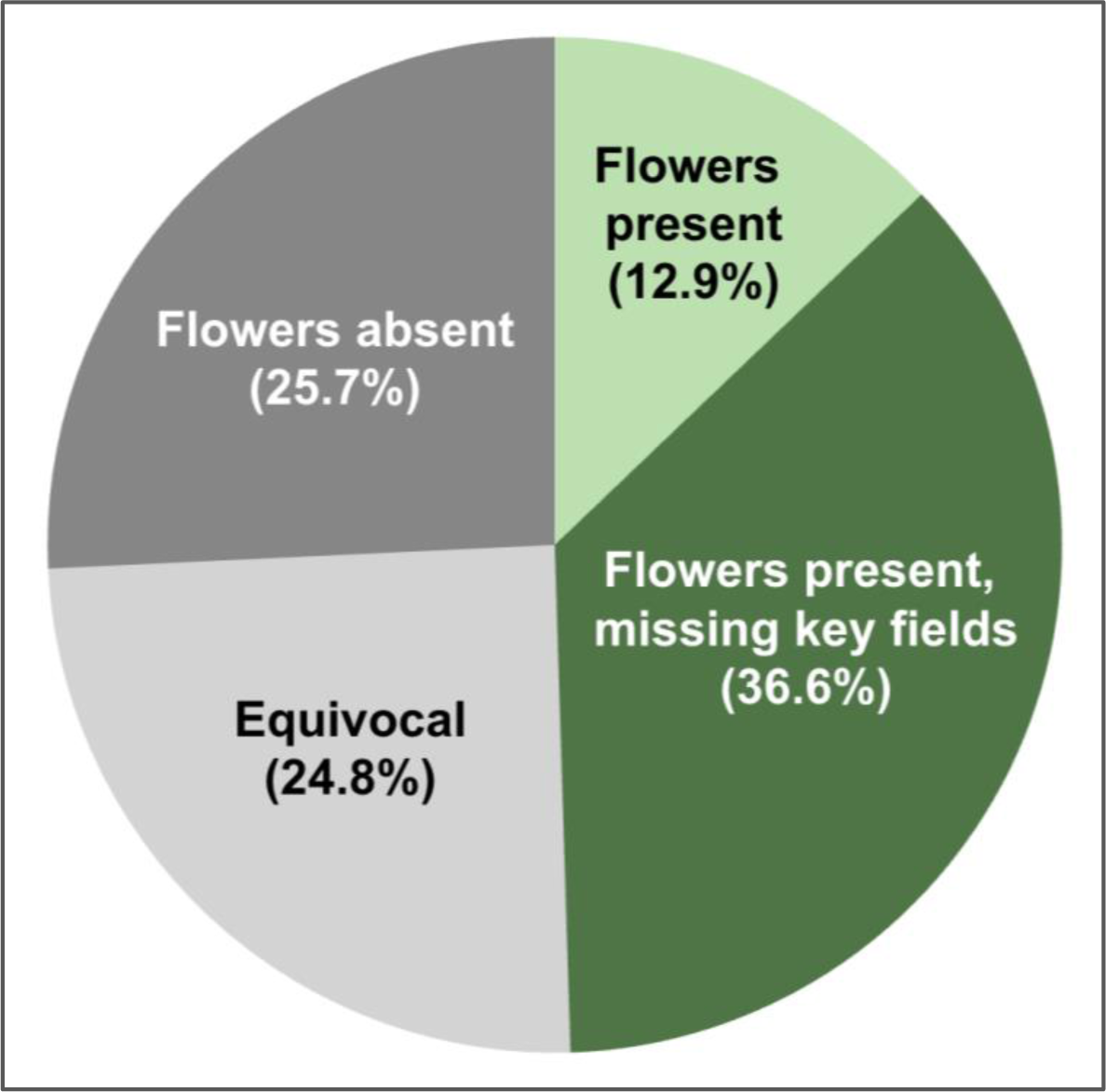
Percent of all records pushed through the ensemble pipeline labeled as “flowers present”, “flowers absent”, and “equivocal”, highlighting that the majority of records labeled as “flowers present” were unusable because they were missing critical date and/or location information. Only records in the “Flowers present” category that had all critical fields were retained, representing 12.9% of all records.

### Data coverage and evaluating specimens missing critical information

The 2,913,35 records annotated as flowers present represented 383 plant families and 8,883 genera (Figure 4). They represent 320 years of collecting, with the earliest record dated in 1501 (Figure 5). We evaluated a small subset of the 8,218,467 records that were missing locality and date information to determine exactly what fields were missing and if those issues could be resolved with more complete digitization of labels. Out of the 100 random specimens that were missing date and latitude/longitude information, 65 had a date (specific day and year) on the specimen label while 35 did not; 85 had a locality that is georeferenceable and localizable to a unit lower than country, while 15 did not. A total of 57 out of 100 had both a georeferenceable specific locality and date information on the label. Only 44 of the 100 records had images retrievable from the original URL, while for 56 records the image URL failed.

**Figure 4.**
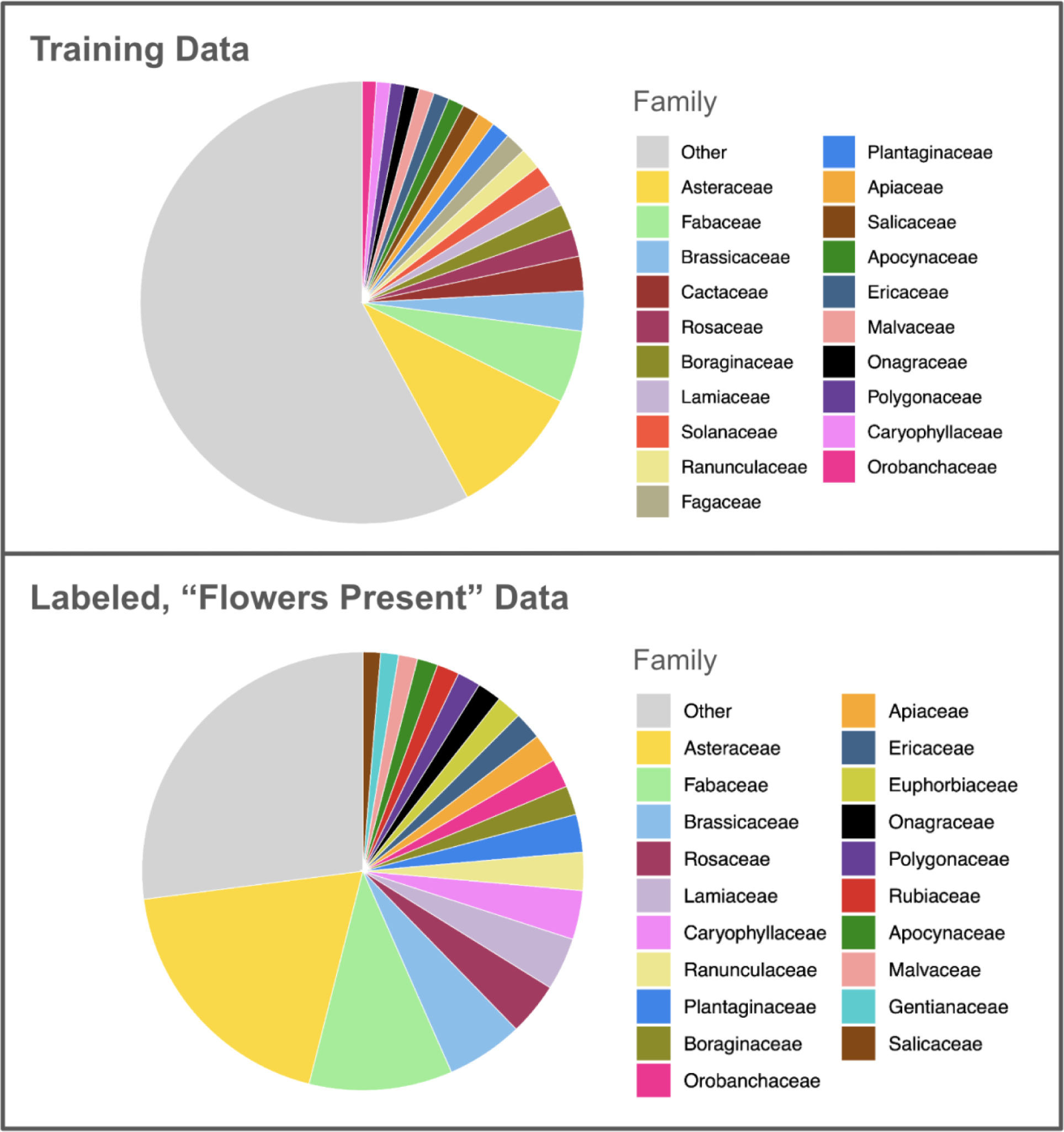
Pie charts showing the top 20 families by number of records in the training dataset (includes training, testing, and validation sets) and in the labeled, present data.

**Figure 5.**
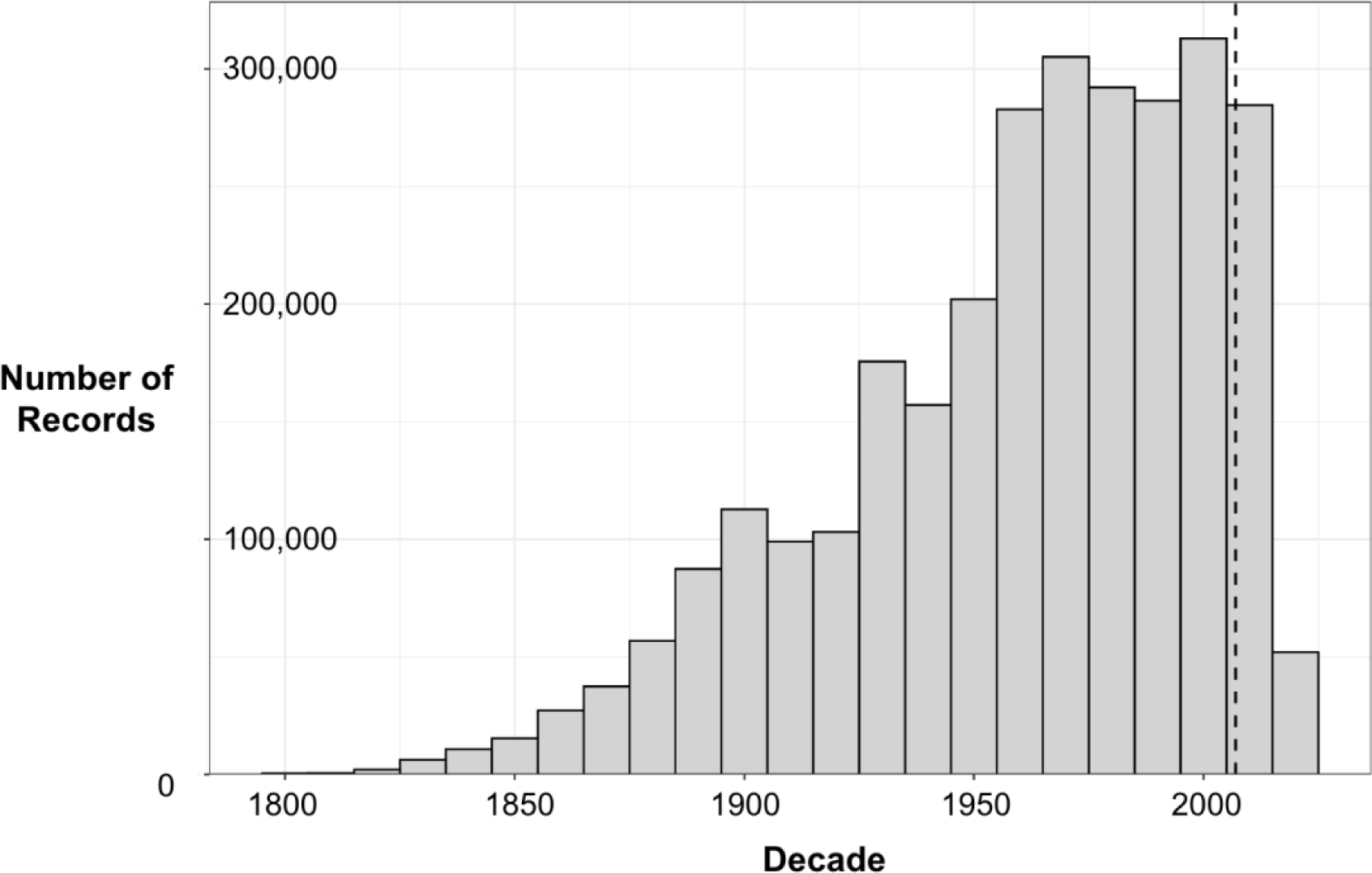
Number of “flowers present” labeled records per decade, beginning at 1800. The dashed line indicates 2007, the year the USA National Phenology Network was founded and the year before iNaturalist began.

## DISCUSSION

While machine-automated annotation is a clear next step in addressing bottlenecks and delivering large quantities of valuable data to the research community, some annotation tasks are harder than others. Further, tasks that are relatively straightforward in one context, such as annotating flowers in photographs of live plants, can be much more challenging in other contexts, such as attempting to do the same for dried, pressed, and aged plants on herbarium sheets. Other groups have produced herbarium data using machine labeling approaches to answer their own research questions about machine learning, certain taxa, or regions, and many showcase reasonably high precision and recall (Goëau et al., 2020; Lorieul et al., 2019; Williamson et al., 2025; Younis et al., 2020). Here, however, our focus was on creating a reliable resource at the broadest possible scale and for the phenology research community as a whole; we were not focused on answering specific research questions. Our first priority was that stakeholders could trust, understand, and easily use these data. A key challenge for us was to develop a set of solutions that we felt met a reasonable bar for trustworthiness, which lead to rigorous data filtering, and to clearly communicate where these solutions may fall short. Here we are not wholly concerned with the size of the data product or any other metric of success. Rather, creating trustworthy data in this context meant carefully vetting the quality of training data, evaluating validation statistics, and being thoughtful about the data structure and documentation made available to stakeholders so they can best use these data and understand their value and limitations.

We also caution that the most appealing solutions for annotating flowers may be difficult at the taxonomic, spatial and temporal scale we attempted here. Our initial hope when generating training data was that we could produce flower annotations utilizing a team of people, likely undergraduate students, with limited botanical training. While this approach might work when focusing on a small subset of plants, such as a genus or family, when scaling to all angiosperms we found it untenable. A vast, complicated diversity of inflorescence morphologies interacts with herbarium sheet taphonomic processes to make it especially challenging to detect flower presence, even for an expert botanist. We argue that the ideal solution for creating high quality annotations is delegating annotation tasks to botanists working on taxonomic groups where they have expertise, but this is also impractical given capacity and logistical constraints. Our solution here was using one expert with ample practice scoring sheets, which has both benefits and drawbacks. One major benefit is that there is likely more consistency and higher quality with this method than a crowdsourced solution. This is especially important in dealing with the many cases where flower presence was uncertain, which was critical for removing certain families and genera from downstream annotation labeling. However, a major drawback in using only one annotator to generate training data is that this method is much less scalable, generating significantly fewer annotated records to use for training. Finally, more focused expertise in certain taxonomic groups may have produced less “unknowns” and higher quality training data for those groups that could lead to inclusion in downstream modeling.

A key step in our process was making thoughtful, informed decisions about removing taxa that we felt were too challenging to score correctly. While annotating training data, the annotator could choose “present” or “absent” options to inform the models, but critically they also had an “unknown” option which they were encouraged to select whenever they could not make a confident determination. Ultimately, the proportion of “unknown” annotations, for individual genera and also families, was what we used to determine which taxa were simply too challenging to score consistently for flower presence. There is likely no perfect cutoff, but we decided that taxa where more than 25% of records were too difficult to label were likely to be problematic for modeling. This decision represents a tradeoff between data quantity and quality. While it is always disappointing to exclude data, we prioritized maintaining stakeholder trust by favoring reliability over volume. We maintain that when attempting to scale challenging tasks such as phenological annotation of herbarium specimens, it is critical to acknowledge both human and technological limits and adjust methods accordingly.

Even with substantial taxonomic filtering, model performance was middling on both held-out test data and expert validated data. Given the challenges with fitting a single model, we opted for an ensemble approach, which gave us more power to interrogate multiple models and arrive at a consensus. Ensembling is particularly useful because it reduces idiosyncratic errors from any single model. We required a two-thirds majority to score presence from the 3 separate models. This strategy required more up-front effort, but ultimately produced a more rigorous labeling workflow. As in other approaches we have published (Dinnage et al., 2025), our interest is in phenophase presence, and the ensembled models are fairly good at avoiding false positives, but we note that the model still produced a substantial number of false absences. Since we explicitly tuned the models to minimize false positives at the cost of increasing false negatives, it means we are likely throwing out many images that have flowers.

The filtering process didn’t end after we completed machine labeling. Out of 11 million records labeled as flowers present, we found that more than 70% lacked a reported latitude/longitude and/or a date. This is consistent with the overall proportion of specimen records on GBIF that have missing coordinates or spatial issues. When we queried GBIF on December 16, 2025, for example, there are 52,254,706 records with associated specimen images for Magnoliopsida and Liliopsida. However, only about 32% of them (16,786,460) have latitude and longitude coordinates and without geospatial issues. This data incompleteness percent is disappointingly high and effectively limits the utility of these data for research, as date and location information is generally required for these data to be research ready (sensu Soltis et al., 2018). The root cause of this issue appears to be due to the publication of “stub records” to GBIF. These stub records may have taxon names but often lack dates and/or georeferences and they generally are from very large collections. For example, the Museum National d’Histoire Naturelle de Paris published 5,031,777 angiosperm records with images and this provider was one where records were often found to be lacking date and geospatial data. Further, we should note that many of the images we annotated are no longer available on GBIF. Out of the 100 we checked, more than half no longer had resolvable images online; most of those also came from publishers who often supply a large corpus of images. These significant issues bring home the need for efforts to expedite full label digitization to avoid these issues of missing metadata since many labels do contain dates and can be georeferenced. Such efforts are needed to support the best use of primary and secondary data (sensu Soberón and Peterson, 2004), such as phenology annotations generated from images. Creating more persistence for images tied to specimens will also better ensure reproducibility and stability when relating specimen images, core specimen metadata and secondary data derived from those images.

Despite setbacks, our ensemble modeling flower annotation pipeline unlocks vast new global phenology data at unprecedented temporal scales. Although in situ phenology data collection networks (Crimmins et al., 2022) and other initiatives to unlock phenology information from sources like iNaturalist (Dinnage et al., 2025) are providing large quantities of contemporary phenology data, data predating these initiatives are less common. They are often in the form of tabular reports that were monitored at one site over many years, although some larger citizen science projects covered multiple sites and years such as the New York Phenology Project (Fuccillo Battle et al., 2022) and Eastern USA Phenology Project (Guralnick et al., 2024) in the middle and late 1800’s. The almost 3 million labeled records that we present here represent a large dataset of historical (pre-2000) phenology now accessible for the research community. However, we note that significant gaps remain in phenology data coverage across space and time, and that continued digitization and efforts to add missing metadata to existing digitized records will be essential for continuing to close these gaps.

All records labeled as flowers present from this effort are available through Phenobase, an open-access datastore of plant phenology information that integrates in situ data from phenology monitoring programs, machine-labeled data from iNaturalist images, and now machine-labeled data from herbarium specimens. Our pipeline produces phenology labels and fields consistent with the Phenobase data structure and Plant Phenology Ontology (Stucky et al., 2018), and uses a controlled vocabulary for other fields, ensuring that data users can search for phenology records at any level of granularity and get consistent results. Future work may include efforts to label herbarium specimens for fruits, however our early explorations found this task to be even more challenging than labeling for flowers, likely requiring greater human effort, and yet more rigorous filtering steps. In sum, despite the challenges, providing information on historical flowering dates across the globe and integrating with other data sources enables new analyses and insights spanning regions and taxa, enabling yet broader and more synthetic questions to be asked across temporal, taxonomic and spatial scales.

## Acknowledgements

The authors would like to thank the many individuals who have contributed to collecting, processing, digitizing, and maintaining herbarium specimens, as this effort would not be possible without their dedication. Phenobase is a team effort and the authors acknowledge team members Carrie Seltzer, Ramona Walls, and Taran Lichtenberger. UFIT Research Computing provided computational resources and support. This work was funded by the National Science Foundation (DBI2223512 to Robert Guralnick and DBI2223508 to Daijiang Li).

## Author Contributions

Robert Guralnick, Daijiang Li and Raphael LaFrance initially conceived of this effort. Erin Grady operationalized most aspects of gathering training data, in consultation with Raphael LaFrance and Robert Guralnick. Raphael LaFrance developed the modeling framework for this effort with help from Russell Dinnage. Raphael LaFrance produced final models and descriptive statistics and Erin Grady led expert model validation. Ellen Denny and John Deck helped ensure that data in the right format could be ingested into Phenobase. Erin Grady and Robert Guralnick archived data. Erin Grady and Robert Guralnick wrote the manuscript with help from Raphael LaFrance, Russell Dinnage and Daijiang Li. All authors contributed to drafts and gave final approval for publication.

## Data Availability Statement

The ensemble data models and a corresponding JSON file with model metadata data are housed on Zenodo (https://doi.org/10.5281/zenodo.17079402). Images used in training, validation, and testing are located here: https://zenodo.org/records/17675089. Code used for this project can be found on github (https://github.com/rafelafrance/phenobase/tree/v1.0.0). Training data and final ensemble output can be found on Zenodo (https://doi.org/10.5281/zenodo.17675089).

## Supporting Information

Additional Supporting Information may be found online in the Supporting Information section at the end of the article.

Appendix S1. List of difficult-to-annotate genera and families removed from training and downstream data.

Appendix S2. Table S1. Validation results for held-out test data.

## Notes

### Competing Interest Statement

The authors have declared no competing interest.

