## Appendix S1 for "Petal to the metal: The slow road to automating large-scale phenology labeling for herbarium specimens"

**Removed genera**

alchemilla

alnus

amaranthus

ambrosia

antennaria

aristida

artemisia

asarum

atriplex

bouteloua

callitriche

carex

ceratophyllum

chamaesyce

chenopodium

craspedia

crassula

cuscuta

cyperus

elodea

eragrostis

freycinetia

galium

hedyosmum

juncus

lechea

ludwigia

mitella

mollugo

muhlenbergia

myriophyllum

najas

nolina

panicum

paronychia

parthenium

peperomia

persicaria

phoradendron

piper

platanus

poa

potamogeton

pyrenacantha

rumex

sporobolus

tillandsia

typha

xerochlamys

**Removed families**

amaranthaceae

araceae

cyperaceae

eriocaulaceae

marantaceae

poaceae

urticaceae
