## Appendix S2 for "Petal to the metal: The slow road to automating large-scale phenology labeling for herbarium specimens"

**Table S1.** Validation on held-out test data. Percentages for detected and undetected represent the accuracy of model annotations as compared to expert annotations in the held-out test set. “Equivocal” values represent the percentage of images classified as equivocal. Shaded calls represent model-annotator agreement for present annotations included in the final dataset. The present, equivocal category is bolded and represents how many present images were “lost” from the dataset.

| **Expert annotation** | **Detected** | **Undetected** | **Equivocal** |
| --- | --- | --- | --- |
| Present | 794 (96.4%) | 67 (13.1%) | **54 (5.9%)** |
| Absent | 30 (3.6%) | 443 (86.9%) | 54 (10.2%) |
